# DRG Explant Model: Elucidating Mechanisms of Oxaliplatin-Induced Peripheral Neuropathy and Identifying Potential Therapeutic Targets

**DOI:** 10.1101/2023.10.05.560580

**Authors:** Junwei Du, Leland C. Sudlow, Igor D. Luzhansky, Mikhail Y. Berezin

**Affiliations:** Mallinckrodt Institute of Radiology, Washington University School of Medicine St. Louis, MO 63110, USA; Institute of Materials Science & Engineering Washington University, St. Louis, MO 63130, USA

## Abstract

Oxaliplatin triggered chemotherapy induced peripheral neuropathy (CIPN) is a common and debilitating side effect of cancer treatment which limits the efficacy of chemotherapy and negatively impacts patients quality of life dramatically. For better understanding the mechanisms of CIPN and screen for potential therapeutic targets, it is critical to have reliable *in vitro* assays that effectively mirror the neuropathy *in vivo*. In this study, we established a dorsal root ganglia (DRG) explant model. This model displayed dose-dependent inhibition of neurite outgrowth in response to oxaliplatin, while oxalic acid exhibited no significant impact on the regrowth of DRG. The robustness of this assay was further demonstrated by the inhibition of OCT2 transporter, which facilitates oxaliplatin accumulation in neurons, fully restoring the neurite regrowth capacity. Using this model, we revealed that oxaliplatin triggered a substantial increase of oxidative stress in DRG. Notably, inhibition of TXNIP with verapamil significantly reduced oxidative stress level. Our results demonstrated the use of DRG explants as an efficient model to study the mechanisms of CIPN and screen for potential treatments.

## INTRODUCTION

Oxaliplatin, a third-generation platinum-based chemotherapy drug, is predominantly used in the treatment of colorectal cancer, with recent research broadening its application to include lung, gastric, ovarian, and prostate cancers (1-4). One of the most common side effect associated with oxaliplatin treatment is peripheral neuropathy, which develops in a ‘stocking and glove’ pattern in patients’ feet and hands (5). Symptoms of oxaliplatin triggered CIPN can appear minutes to hours after treatment, and are likely to persist and accumulate with repeating dosages, resulting in chronic symptoms lasting months, or even years later. CIPN affects patients’ every aspect of daily life, compelling clinicians to lower the dose of cancer chemotherapy agents and consequently decreasing the therapeutic efficacy (6, 7).

The mechanisms by which oxaliplatin exerts its anti-tumor effects are largely due to its ability to damage DNA and generate excess reactive oxygen species (ROS) that lead to apoptosis in cancer cells (8, 9). While these mechanisms are crucial for the drug’s anti-cancer activity, they can also have unintended consequences on healthy cells (10), including those in the peripheral nervous system (PNS). PNS especially dorsal root ganglia (DRG) appear to be highly susceptible to the chemo-induced oxidative stress as a result of lacking vascular barriers and an absence of lymphatic drainage (11). Peripheral neurons are also highly vulnerable to oxidative stress (10, 12) due to their high metabolic rate, long axonal structures, and the presence of supporting myelin and satellite glial cells (SGCs), which can itself be damaged by ROS. Once the neurons are damaged, they trigger a cascade of events that may result in symptoms like tingling, cold/hot allodynia, numbness, and eventually chronic pain, the hallmarks of CIPN.

Significant effort has been aimed studying how to mitigate CIPN side effects, including the use of antioxidants and other neuroprotective agents, without compromising the drug’s anti-cancer efficacy (13-15). However, clinical trials targeting ROS had not yet demonstrated significant relief from the symptoms. This discrepancy highlights the complexities involved in CIPN and raises questions about the effectiveness of simply suppressing ROS with antioxidants as a therapeutic strategy that so far only provides mild to moderate pain relief (16, 17).

Finding better treatments for CIPN may require different types of assay. Most of the published techniques used to understand the mechanism of CIPN rely on either cell cultures or live animals. Both of these traditional models have substantial limitations that impede the discovery of effective therapies. For example, neuronal cell cultures that are often used to study oxaliplatin-induced ROS (18, 19) are too simplified as a model and do not replicate the complex microenvironment found in vivo. These models focused on isolated neurons do not take into account other types of cells such as SGCs, Schwann cells, or immune cells, which play significant roles in the pathogenesis of CIPN. Neuronal cultures also have limited lifespans, restricting the length of time over which experiments can be conducted, which may not be representative of chronic conditions like CIPN. Animal models, such as based on wild type mice treated with chemotherapy drugs introduce a host of systemic variables such as metabolic changes (20), immune responses, and changes in the blood composition, which can confound results and make it challenging to isolate the specific impacts of a drug on CIPN. In addition, live animal models have low throughput and require a lengthy period to develop chronic CIPN.

Given these limitations, new assay types are needed for CIPN research that combine the throughput of the neuronal culture while having substantial relevant complexity such as in live animals. We envisioned that dorsal root ganglia (DRG) that houses clusters of sensory neuron somata located along the spinal cord. The DRG play a key role in transmitting sensory information and pain signals from the periphery to the central nervous system and being isolated might be used as alternative CIPN assay. These isolated DRG ex vivo models are increasingly being used for studying a variety of neurological diseases and for screening potential drugs (21). Unlike simplified cell culture models, DRG models retain the native arrangement and diversity of cell types, including neurons, SGC, and other supporting cells, providing a more biologically relevant environment. DRGs are also physiologically relevant and represent a natural and intact microenvironment that preserves the native cellular interactions and signaling pathways found *in vivo*. Compared to the live animal models, where systemic factors like immune response and metabolic rate can influence outcomes, extracted DRG models help eliminate these confounding variables and isolate the effect of the drug on sensory neurons.

We hypothesized that the neurotoxicity of oxaliplatin, and potentially other drugs that cause ROS neurotoxicity, may be quantified by using a novel in vitro model of extracted DRG. This assay can be also used for studying pathways and testing new molecules as potential drugs to prevent or treat CIPN. Specifically we were interested in understanding the role of TXNIP mediated ROS – inflammatory pathway as a trigger to CIPN. To test this hypothesis, we extracted DRG from mice, measured neurite outgrowth, evaluated oxidative stress and tested TXNIP inhibitors, aiming to understand the pathology of CIPN and to identify therapeutic targets.

## METHODS

### Animals

8-12 weeks old, male B57CL/6 male mice from Charles River Laboratory were used in this study. Animals were housed with free access to water and food and maintained on a 12:12 hour light/dark cycle with controlled temperature and humidity. All animal experiments were conducted in compliance with Washington University Institutional Animal Studies Committee and NIH guidelines.

### Extraction and DRG preparation

Thoracic and lumbar DRGs from adult (8-12 weeks) healthy male C57BL/6 mice were aseptically removed from levels L1-L6 and transferred into ice-cold serum-free media (SFM), which was a customized DMEM-based media (ThermoFisher Scientific, cat # A4192101) containing 1% BSA (Millipore Sigma, cat # A7906), 50 µg/mL vitamin C (Millipore Sigma, cat # A4544), 3 g/L extra glucose (Millipore Sigma, cat # G8270), 1% L-glutamine (ThermoFisher, cat # 25030149), 1% insulin-transferrin-selenium (ThermoFisher, cat # 51300044) and 1% penicillin-streptomycin (ThermoFisher, cat # 15140122). The DRGs were trimmed to remove any fibers connecting to the ganglia. A stock solution of Matrigel-SFM mixture (1:1 vol) was prepared by mixing growth-factor-reduced Matrigel (Corning, cat # 354230) with SFM in a sterile Eppendorf tube until homogenized and kept on ice. Matrigel-SFM mixture (20 µ L) was placed in each well of a 12 well plate (Corning, cat # 3513). DRGs were placed individually in each well into the Matrigel-SFM mixture and incubated in 5% CO_2_ at 37□ °C for 50□ min. Finally, 1.5 mL pre-warmed SFM was added to each well and DRGs were incubated at 37□°C for further use.

### Fluorescent imaging

Imaging was performed with Zeiss Cell Discover 7 confocal microscope. All images were captured using a 5x objective. The pixel time was set at 0.52 μs. Three laser wavelengths were used for immunochemistry staining:561 nm for Alexa Fluor 555 with a detection wavelength range of 550-700 nm, 488 nm for Alexa Fluor 488 with a detection wavelength range of 400-550 nm, and 405 nm for DAPI, also with a detection range of 400-550 nm. For ROS imaging, laser wavelength was 488 nm and detection range was 490-700 nm.

The ROS level in the DRG was determined by dividing the fluorescence intensity within the DRG region of interest (ROI) by that of the ROI outside the DRG. The calculated value was then normalized to the control values at the time zero.

### DRG outgrowth assay

Freshly excised isolated DRGs were placed on a Matrigel-SFM mixture (see above) at the bottom of individual wells of a 12-well plate and placed in an incubator with 5% CO_2_ at 37°C. SFM media was replaced on the 3^rd^ day. The outgrowth assay for the control untreated DRG was determined within 5 days. The DRG were periodically imaged with a bright field inverted microscope to monitor the neurite outgrowth (Olympus CKX53). Images of DRG neurites outgrowth were analyzed using the Sholl Analysis plugin in ImageJ. Briefly, the explant DRG was excluded by a circle enveloping it, and rings were created around it in 5 µm increments. The number of intersections that neurites crossing each ring was counted and the neurite occupied area was represented by the total number of intersections of all rings.

### Treatment of DRG with oxaliplatin

To investigate the effects of oxaliplatin on the DRGs regrowth, the media for the oxaliplatin-treatment DRG’s was replaced 48h after plating with SFM mixed with different doses of oxaliplatin (in sterile water) and incubated for an additional 72 h.

### Immunochemistry of DRG

After treatment with oxaliplatin or other drugs, DRGs were fixed in 10% neutral buffered formalin (NBF) for 30 min at room temperature. Following additional incubation with a 1X PBS buffer containing 0.1% Triton-X (PBST), DRGs were blocked with 10% donkey serum (Millipore Sigma, cat # D9663) in 0.1% PBST. DRG explants were then incubated with the primary antibody against β-III tubulin (Millipore Sigma, cat # MAB1637, 1:500) and calcitonin gene-related peptide (CGRP) (ThermoFisher, cat # PA185250, 1:500) overnight at 4°C, followed by incubation with Alexa Fluor 555 (ThermoFisher, cat # A31570, 1:1000) and 488 (ThermoFisher, cat # A32814, 1:1000)-conjugated secondary antibody for 1h at room temperature. Nuclei were stained with DAPI (ThermoFisher, cat # 62248) for 20 min. The entire DRGs were scanned with Zeiss Cell Discover 7 confocal microscope under 5x objective to generate 16 images that were stitched together to capture the whole DRG and surrounding axons.

### DRG ROS imaging

CM-H_2_DCFDA (ThermoFisher, cat #C6827) or CellRox Green (ThermoFisher, cat #C10444) dissolved in DMSO were added to pre-warmed to 37°C HBSS buffer at a final concentration of 10 µM. DRGs were then incubated with the dye at 37°C for 40 min in the incubator. Then, the buffer was removed, and DRGs were washed with HBSS three times and recovered in the SFM for 10 min. Fluorescence of untreated DRGs were determined as a baseline and recorded periodically within 120 min. To investigate the effects of oxaliplatin and OCT2 inhibitor on ROS, DRGs were exposed to 100 μM oxaliplatin, mixtures of 100 μM oxaliplatin and 1000 μM cimetidine (Millipore Sigma, cat # C4522) in 5% DMSO, or 5% DMSO. For testing effects of TXNIP inhibition on ROS, DRGs were treated with 100 μM oxaliplatin, 100 μM oxaliplatin combined with 1 μM verapamil (Millipore Sigma, cat # V4629), and sterile water. Fluorescence intensity was monitored every 10 min with Zeiss Cell Discover 7 confocal microscope for 120 min under 5x objective with no stitching to capture the whole DRG.

### Statistics and data analysis

The values were then processed in Prism 9 for statistical analysis using one-way ANOVA test with multiple comparisons. Data were expressed in mean ± SD.

## RESULTS

### Growth media induces stable and uniform distribution of neurite outgrowth

DRG neurons inherently possess a distinct regenerative capacity. When exposed to an appropriate environment or substrate, they are naturally programmed to extend axons in the process known as outgrowth. To facilitate the complete DRG outgrowth, we used Matrigel as an scaffold to which the DRG could attach. Matrigel is a viscous liquid derived from Engelbreth-Holm-Swarm mouse sarcoma that primarily comprises extracellular matrix (ECM). To minimize the role of growth factors, we selected Matrigel with low level of growth hormones. Isolated DRG cultured on Matrigel benefit from the matrix’s unique properties that promote axon outgrowth (22). Occasionally, a DRG failed to attach to the Matrigel and consequently failed to show any neurite outgrowth (data not shown). By offering a scaffold, Matrigel replicates the in vivo extracellular environment (23), supporting both cell adhesion, survival and growth.

Under our standard growth media conditions (Matrigel-SFM, 37°C, 5% CO_2_) DRGs showed stable and evenly-distributed neurite outgrowth. Bright-field images of DRG (**Supplemental, Figure S1)** outgrowth were difficult to quantify due to the lack of contrast between the background and small fibers in bright field views. To overcome this limitation, we used fluorescent labeling against the neuronal marker β-III tubulin and employed high-resolution fluorescence microscopy (see Methods) with stitching capabilities. This imaging method allowed us to capture the entire DRG outgrowth with single neurite resolution. High-resolution fluorescent imaging data were further quantified using Sholl analysis, a technique commonly used to quantify regrowth pattern in the DRG (24, 25). This computational technique provided a systematic way to measure the neurite extensions’ complexity and extent, as illustrated in **Error! Reference source not found.A**. The method involves placing concentric circles at regular 5 um intervals around the entire DRG (26). The number of times neurites intersect these circles is counted, and the data are represented as a Sholl profile (**Error! Reference source not found.B**). On this profile, the x-axis indicates the distance from the soma (i.e., the radius of the circles), and the y-axis shows the number of intersections at each radius. The example of Sholl analysis for the DRG is shown in Error! Reference source not found.**A-B**.

### Oxaliplatin induces decreases in DRG neurite outgrowth in a dose-dependent pattern

The addition of oxaliplatin to the SFM culture media had a profound and dose dependent effect on the DRG outgrowth as shown in **Figure 1**. The media with 50 μM level of oxaliplatin yielded inhibition of almost half (48%) of the neurite outgrowth, and 100 μM oxaliplatin inhibited more than 70% of neurites compared to the control group during antibody staining against bIII-tubulin. CGRP(+) neurons that are associated with the transmission and modulation of pain signals, both nociceptive (responsive to potentially damaging stimuli) and neuropathic (resulting from damage to the nervous system) (27, 28) were similarly affected (**Figure 1E-F**). The ratio of CGRP(+)/β-III tubulin(+) neurons in the regrown DRG showed no statistical difference between control and oxaliplatin-treated groups (**Supplemental**,, **Figure S2**), suggesting lack of specificity during the process of neuronal regrowth in the DRG in the presence of oxaliplatin.

**Figure 1.**
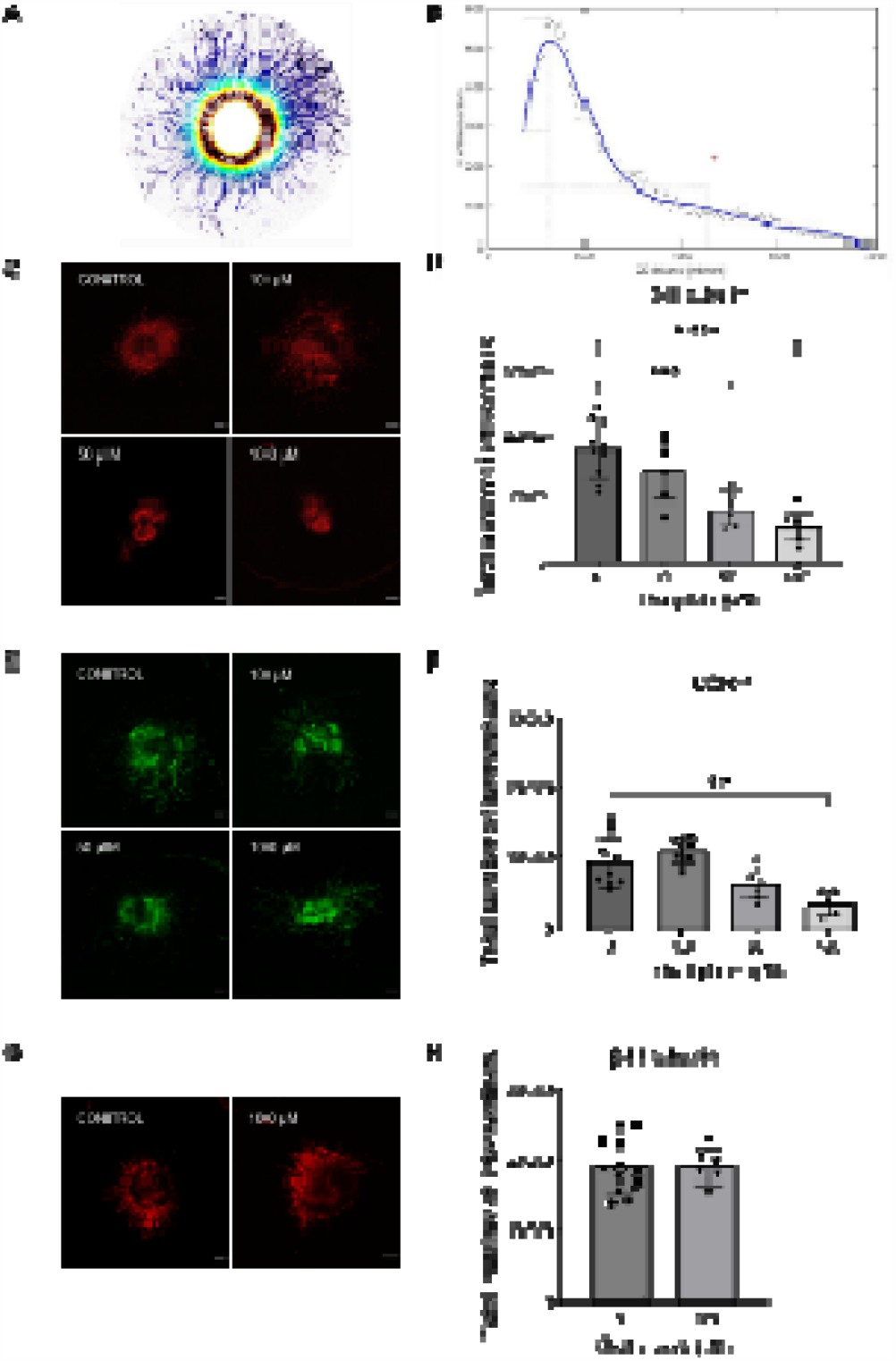
Oxaliplatin induced dose-dependent reduction of neurite outgrowth in DRG explants. (**A-B**) Neurit outgrowth was quantified with Sholl Analysis. Representative images of Sholl mask (**A**) and Sholl profile (**B**) displayed the counts of intersections per sampling shell against distance from center (generated in ImageJ). Neurite occupied area can be represented by total number of intersections. (**C**) β-III tubulin staining of DRG explant after 72h treatment with 0, 10, 50 and 100 μM oxaliplatin. (**D**) Oxaliplatin suppressed neurite outgrowth in a dose-dependent manner. Statistical significance was detected at concentration of 50 and 100 μM oxaliplatin. (**E**) CGRP staining of DRG explant after 72h oxaliplatin treatment. (**F**) Oxaliplatin leads to decrease of CGRP(+) neurites in DRG. (**G**) β-III tubuli staining of DRG after 0 and 100 μM oxalic acid treatment. (**H**) Oxalic acid did not affect the neurite outgrowth of DRG explants. Scale bar: 200 μm. Zeiss Cell Discover 7, obj 5X, stitched images. Data are presented in mean ± SD. **P<0.01, ***P<0.001, ****P<0.0001.

### Oxalate does not affect DRG regrowth

One subject of debate is the role of oxalate in the development of CIPN. Oxalate is not only a key component of oxaliplatin’s structure — serving as a ligand that chelates Pt^2+^ — but is also a major metabolite during oxaliplatin treatment. Several studies have suggested that the strong chelating properties of oxalates could bind Ca^2+^ ions in neurons, thereby affecting the function of voltage-dependent sodium channels (29, 30). This alterative model has been proposed as a primary factor in the acute neurotoxicity induced by oxaliplatin. To examine the role of oxalate, we performed DRG outgrowth assays in its presence at concentrations equivalent to those of oxaliplatin. We observed no statistically significant differences between DRGs treated with oxalate at comparable doses of oxaliplatin (100 μM) and control ones (**Figure 1G-H**). These findings suggest that oxalate alone has a minimal effect on DRG neurite outgrowth, challenging the hypothesis that oxalate is a key factor in CIPN (7, 31-33).

### Inhibition of OCT2 preserves DRG neurite outgrowth

Having demonstrated the negative impact of oxaliplatin on DRG neurite regrowth, we further validated the assay’s utility by inhibiting oxaliplatin uptake into the SGCs. Recent studies in mouse models have shown that the organic cation transporter 2 (OCT2) mediates the uptake of oxaliplatin into SGCs, initiating neuropathy symptoms. Blocking this transporter with an OCT2 inhibitor cimetidine (34) has been previously shown to minimize these symptoms (35, 36). To investigate whether OCT2 inhibition could protect neurite outgrowth in the presence of oxaliplatin, DRG explants were co-cultured with a high dose of oxaliplatin (100 μM) and varying concentrations of cimetidine. We observed a significant increase in the area occupied by neurites in the DRG explants treated with both cimetidine and oxaliplatin compared to those treated solely with oxaliplatin (**Figure 2**). The protective effects became statistically significant at a cimetidine concentration of around 100 μM, with an EC50 value of approximately 159.6 μM. These results are in line with the published data on the neuroprotective properties of cimetidine in oxaliplatin-treated mice, further validating the assay’s relevance to animal studies.

**Figure 2.**
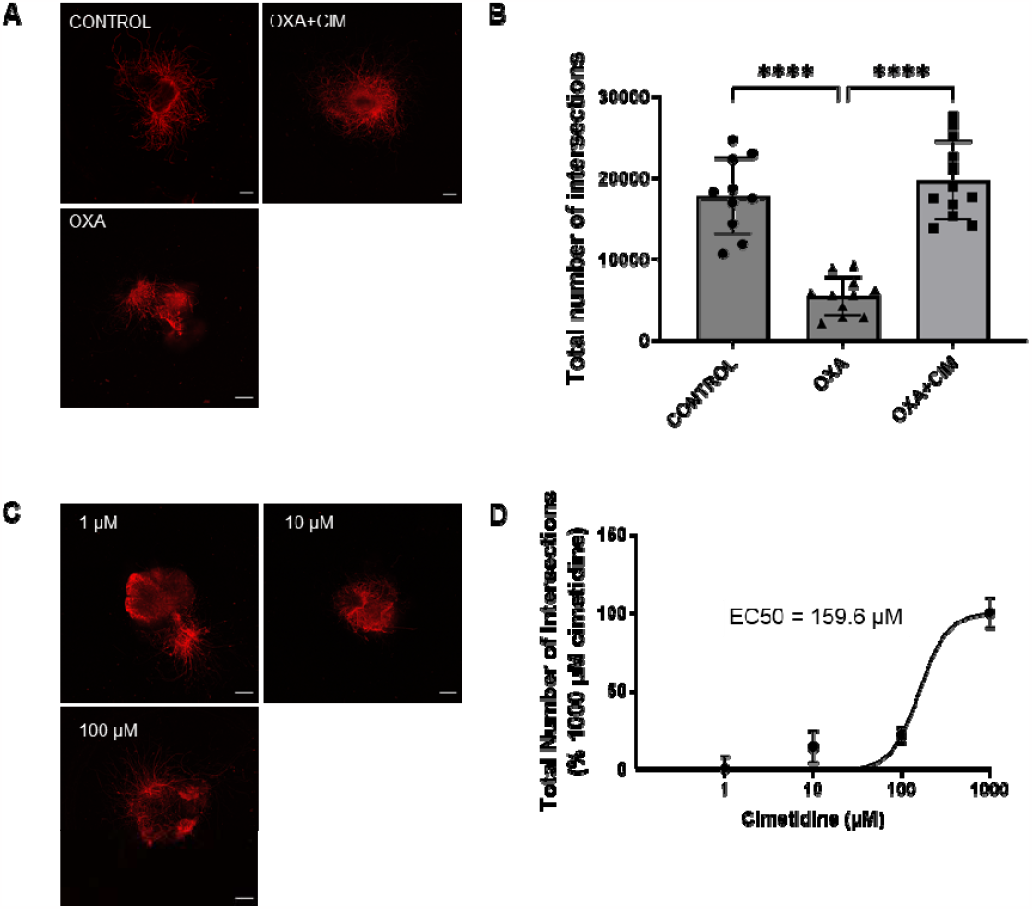
Inhibition of OCT2 preserve neurite outgrowth under oxaliplatin treatment. (A) β-III tubulin staining of DRG explants under DMSO (control), 1000 μM cimetidine and 100 μM oxaliplatin, and 100 μM oxaliplatin treatment for 72h. (B) 1000 μM cimetidine led to significant improvement of neurite outgrowth under oxaliplatin. There was no significance between DRGs with no drug treatment and those treated with 100 μM oxaliplatin in presence of 1000 μM cimetidine. (C) β-III tubulin staining of DRGs co-cultured with 100 μM oxaliplatin and varying concentration of cimetidine: 1 μM, 10 μM, and 100 μM. (D) Cimetidine improved neurite outgrowth in a dose-dependent pattern. Th estimated concentration associated with 50% neurite regrowth (EC50) is around 159.6 μM. Scale bar: 200 μm. Zeiss Cell Discover 7, obj 5X, 16 stitched images. Data are presented in mean ± SD. ****P<0.0001.cimetidine: 1 μM, 1 μM, and 100 μM. (**D**) Cimetidine improved neurite outgrowth in a dose-dependent pattern. The estimated concentration associated with 50% neurite regrowth (EC50) is around 159.6 μM. Scale bar: 200 μm. Zeiss Cell Discover 7, obj 5X, 16 stitched images. Data are presented in mean ± SD. ****P<0.0001.

### Oxaliplatin leads to oxidative stress in the DRG explant

While neurite outgrowth assay is an established as an effective assay for assessing oxaliplatin toxicity, it is primarily based on measuring morphological changes. Herein, we extended the use of the DRG explant model to explore the response to oxidative stress. A recent study demonstrated that oxaliplatin could trigger an increase in reactive oxygen species (ROS) production within DRG neurons (18). To investigate whether the same effect can be seen in the DRG, we added activatable fluorescent ROS sensors CM-H_2_DCFDA or CellRox into the media.

CM-H_2_DCFDA is a widely used, cell-permeable probe designed to measure intracellular levels of ROS. Initially non-fluorescent, it easily crosses the cell membrane and becomes trapped inside the cell after esterase enzymes remove its acetate groups. This modified, non-fluorescent charged form, known as H_2_DCF, reacts with ROS in the cell to produce a highly fluorescent compound dichlorofluorescein. The level of ROS can be thus quantified from the intesity of the fluorescence signal using fluoresence microscopy. CellRox Green is a newer generation of ROS probes with the same mechanism of action but with higher photostability according to the manufacturer.

In this assay, the DRG was extracted, placed in a well as per the Extraction and DRG preparation methods section, and allowed to regrow for two days, followed by ROS dye treatment. Fluorescence intensity of the dye in the presence of DRG was rather low and slightly decreased over the duration of the experiment (120 min) when normalized to the background and to the intensity right after addition of the dye (**Figure 3**). In contrast, addition of oxaliplatin substantially increased the fluorescence intensity of the DRG. We observed steady increase of ROS relative to the background when the DRG was exposed to 100 μM oxaliplatin for 120 minutes. Furthermore, the inhibition of OCT2 via cimetidine almost completely prevented the oxaliplatin-induced oxidative stress with the average normalized intensity in the DRG very similar to the control DRG with no oxaliplatin added. These findings suggested the DRG explant as a viable model for investigating the role of ROS in oxaliplatin-mediated DRG tissue damage.

**Figure 3.**
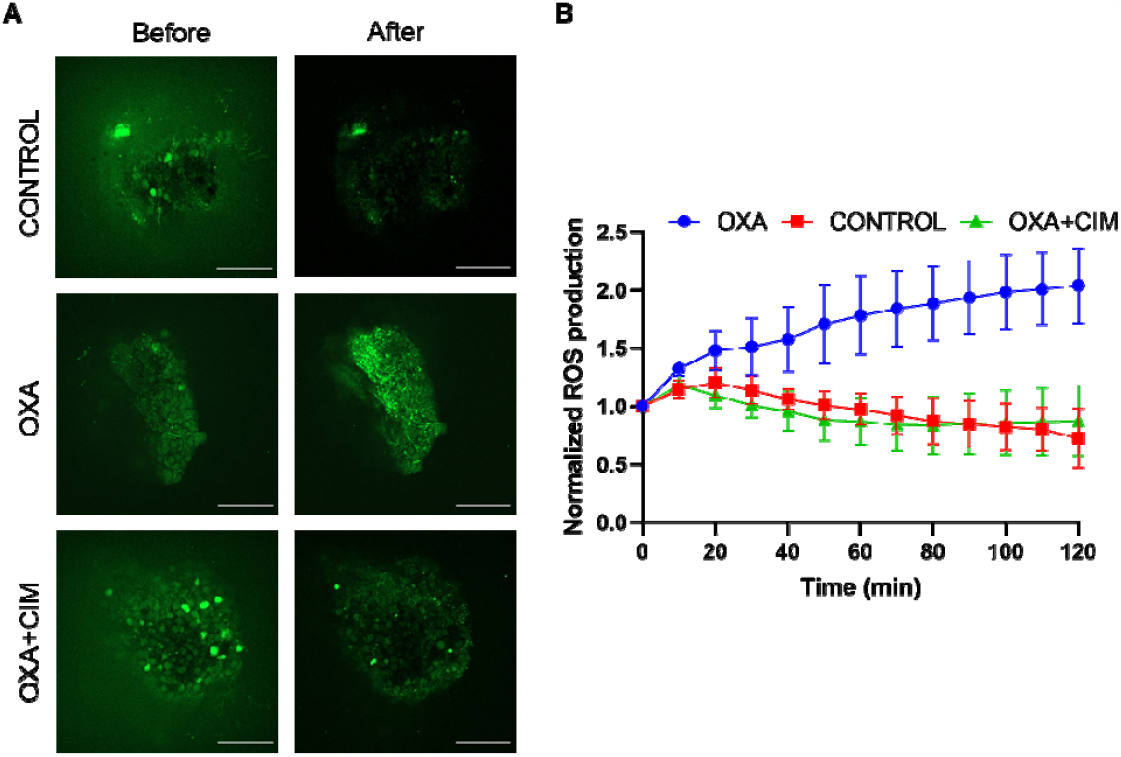
Oxaliplatin led to prolonged increasing of oxidative stress in DRG. (A) Representative CM-H2DCFDA images of DRG explants treated with DMSO, oxaliplatin, and cimetidine (1000 μM) with oxaliplatin (100 μM) at two timepoints: immediately after drug loading (before, time zero), and 120 min after. (B) Quantification of CM-H2DCFDA fluorescence signal revealed oxaliplatin induced increased oxidative stress throughout experimental period. Inhibitio of OCT2 by cimetidine prevented the burst of ROS. Data from the DRG ROI were divided to the intensity of the solution ROI and then normalized to the time point 0. Scale bar: 200 μM. Data are presented in mean ± SD. N=3. Zeiss Cell Discover 7, obj 5X.

### Inhibition of TXNIP through verapamil decreases ROS level induced by oxaliplatin

Encouraged by the establishment of both neurite outgrowth and ROS assay as metrics of tissue damage and oxidative stress, we evaluated the potential neuroprotective effects of inhibiting TXNIP with Verapamil. Verapamil, a clinically approved calcium channel inhibitor, has been previously shown to attenuate neuropathy induced by diabetes through blocking of TXNIP – an enzyme that is controlling ROS in mitochondria and cytosol in mitochondria and cytosol by interacting and inhibiting thioredoxin (37) (see the Discussion). For that, DRG explants were cultured in presence of a high level of oxaliplatin and variable dosages of verapamil. We found 1 μM verapamil resulted in an approximately 70% enhancement in neurite outgrowth compared to the DRGs treated with oxaliplatin (**Figure 4A-B**) with almost complete suppression of the ROS (**Figure 4C-D**).

**Figure 4.**
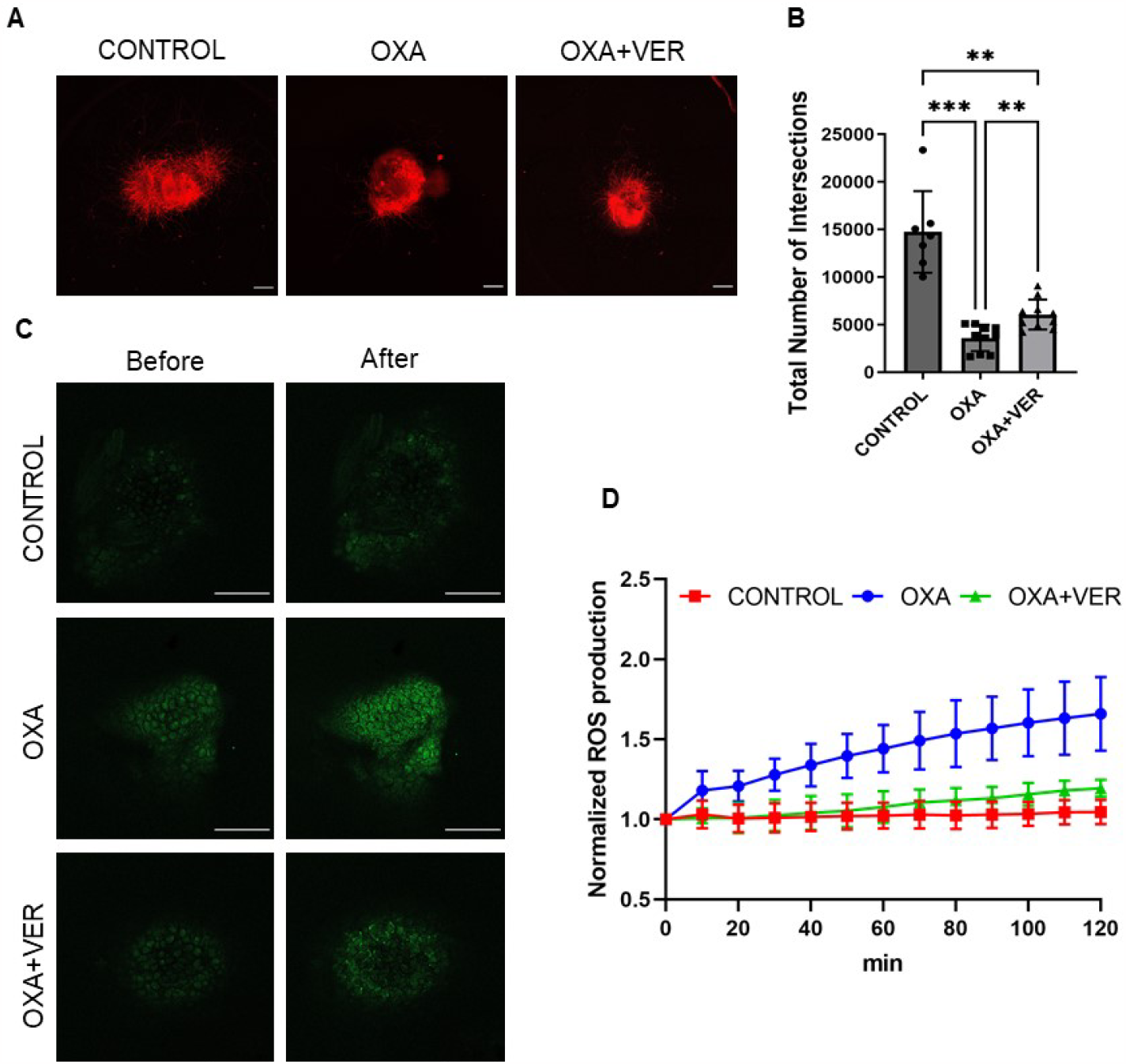
Inhibition of TXNIP through verapamil restore neurite outgrowth capacity and decreases ROS level induced by oxaliplatin. (**A**) Representative images and quantification of total number of intersections of DRG under treatment of vehicle (control), 100 μM oxaliplatin (OXA), and 100 μM oxaliplatin combined with 1 μM verapamil (OXA+VER). (**B**) quantification of total number of intersections of DRG under above treatments. Verapamil resulted in an approximately 70% enhancement in neurite outgrowth compared to the oxaliplatin-only DRGs. (**C**) Representative images of 2-hour short-term ROS detection for DRG explants treated with DMSO (control), oxaliplatin (100 μM), and oxaliplatin (100 μM) combined with verapamil (1 μM) at two timepoints: right after drug loading (time zero), and 120 min. The ROS level in the DRG was determined by dividing the fluorescence intensity within the DRG ROI by that of the ROI outside the DRG. This value was then normalized to the baseline (time 0). (**D**) Quantification of flourescence signal throughout experimental period. Verapamil prevented ROS burst in DRG under oxaliplatin treatment. N=3-4. CellRox Green was used for ROS detection. Scale bar: 100 μm. Data are expressed as the mean□±□SD. **P<0.01, ***P<0.001, ****P<0.0001.

Verapamil was unable to completely restore the DRG to the oxaliplatin at higher dosage, possibly because of its own potential neurotoxic effect (38) (**Supplemental, Figure S3**). Moreover, DRGs co-cultured with verapamil and oxaliplatin demonstrated significantly lower ROS (38). Moreover, DRGs co-cultured with verapamil and oxaliplatin demonstrated significantly lower fluorescence intensity compared to oxaliplatin-only DRGs pointing to the protection effect of verapamil on the DRG (**Figure 4C-D**). These data indicate while verapamil was not able to entirely block oxaliplatin anti-regrowth effect, it substantially mitigated oxaliplatin induced oxidative stress suggesting targeting TXNIP might offer a promising approach to attenuate the inflammatory effects of oxaliplatin on DRG.

## DISCUSSION

### Platinum Drug Impact on DRG

Existing in vivo models require large number of animals and long duration procedures which limits their application in the examination of new mechanisms in early-stage drug discovery and unables high throughput screening (39, 40). In response to these challenges, several in vitro models have been established to emulate CIPN and to screen for possible targets more efficiently. One of the most common in vitro methods relies on neurons derived from either animal embryonic DRGs or human stem cells (41-44). Other single-neuron model rely on neuronal cell lines such as PC-12 and SH-SY5Y (45, 46). While these published studies provide valuable insights into the general neurotoxicity of chemotherapy drugs, models using a single neuronal cell type are often overly simplified and may not adequately capture the complex responses of DRGs to stimuli in their natural environment.

To address this problem, we established a quantitative assay based on DRG explants obtained from adult animals. The usage of adult DRG explants reflects a more realistic effects of oxaliplatin as it retains the intact cell network and extracellular microenvironment that play an important role in neuronal functions (47). DRG explant also offers(47). DRG explants also offer a diversity in cell type, since DRG consists of not only sensory neurons, but also non-neuronal supporting cells including SGCs, immune cells, and endothelial cells (48-50). Although sensory neurons are conventionally considered primary targets in CIPN, emerging evidence indicate alterations in the non-neuronal cells contribute to the development of neuropathic pain (51-56). This underscores the value of our DRG explant model, which effectively recapitulates the multicellular nature of DRG and its response to CIPN.

Oxaliplatin and related platinum drugs have been observed to target the DRG and sensory nerves without affecting motor nerves (57). The drugs accumulate in many neuronal organs, such as DRG, spinal cord, brain, peripheral nerves with the DRG been identified as having the highest accumulation of platinum (58). The unique accumulation of oxaliplatin in the DRG is intriguing. An accepted theory attributes this preference to the DRG’s differential vascular characteristics, distinguishing it from the brain and extended peripheral neurons. Unlike the brain and peripheral nerves, the DRG lacks a robust nerve-blood barrier (11), facilitating the drug’s penetration from the bloodstream into neurons and supporting cells. Functionally, the absence of the barrier allows the DRG rapidly respond to stimuli. However, this accessibility also make the DRG vulnerable to toxins, particularly heavy metals like platinum, which can linger within the DRG for extended durations.

The damage of oxaliplatin to the DRG neurons might occur through several pathways. Oxaliplatin’s primary action in neuronal cells mirrors its anti-cancer effect—by forming DNA adducts and disrupting DNA replication. When sensory neurons are damaged, for example by heavy metals, SGCs proliferate to support the neurons (59). Disturbance of this process stops neurons from getting needed supply, which further damages neurons. Another effect comes from the oxaliplatin induced oxidative stress, which lead to cellular damage. Oxidative stress was the focus of this work.

We use the extracted DRG explant model to evaluate the effect of oxaliplatin, measure the oxidative stress and identified potential treatment. We observed that oxaliplatin significantly reduced neurite outgrowth of DRG explants in vitro in a dose-dependent manner characterized as neurite occupied area. In vivo, the growth of DRG neurons is carefully regulated to prevent uncontrolled proliferation unless they encounter damage. During the damage of the nerve, the nucleus of peripheral neurons provide instructions that enable the neurons to regrow – the trait that makes peripheral neurons unique as shown in **Figure 1**. In the presence of toxic elements, such as oxaliplatin, the regrowth can be slowed or completely stopped. The quantitative aspect of regrowth (area of regrowth, total length of neurites, maximum length of axons) can be used as a sensitive quantitative assay to evaluate the toxicity and identify the new potential treatment.

Almost complete (ca. 90%) reduction of the normal regrowth of DRG was achieved with 100 μM level of oxaliplatin in the regrowth after 72 h while and a significant decrease was noticed at 50 mM of oxaliplatin. When DRG were treated with lower (10 μM) concentration of oxaliplatin, no statistically significant change were detected. These findings are in line with the DRG neuronal cell culture studies, in which reduced cell viability was only observed with concentrations of oxaliplatin higher than 10 µM (18, 41).

The reduction of regrowth apparently was a result of the intact oxaliplatin molecule or its product of hydrolysis, and not due to free oxalate. It has been suggested earlier that dissociated oxalate ligand, which is usually displaced during the activation of the drug, would mediate the acute neuropathy through chelating of Ca^2+^ and Mg^2+^ ions in the nerve tissue (30, 60). In our assay, oxalate seems did not affect the regrowth. A high concentration of oxalate in the DRG media, equivalent to the oxaliplatin level (100 mM), did not cause any changes in the outgrowth pattern compared to the control oxalic free media (**Figure 1**). Although we have not tested whether free, oxalates can penetrate DRG cells, our finding may indicate that oxalate might not the primary driver behind the chronic neurotoxicity induced by oxaliplatin.

### Inhibition of OCT2 restores DRG outgrowth in the presence of oxaliplatin

The accumulation of oxaliplatin in the DRG cells has been shown to be facilitated by OCT2 (35), a transport protein that is expressed in both primary neurons and SGCs (61). SGCs with a higher expression level of OCT2, apparently absorb more oxaliplatin than that the neurons do. (61). It has been observed that blocking OCT2 with different blockers, such as dasatinib (35) and cimetidine (62), suppresses the buildup of oxaliplatin in the DRG. The SGCs provide the structural support to neurons and regulate substance trafficking to and from the primary neurons (63, 64). It has been observed that blocking OCT2 with several blockers, such as dasatinib (35) and cimetidine (62), suppresses the buildup of oxaliplatin in the DRG. Using the outgrowth assay, we have observed that blocking OCT2 with cimetidine -even amidst high concentrations of oxaliplatin -completely restores the outgrowth of DRG to almost a normal level (**Error! Reference source not found.Figure 2**). The measured EC50 = 159.6 uM was relatively high requiring large doses of cimetidine (almost 10 times compare to oxaliplatin) to restore the DRG regrowth, suggesting the need for more effective inhibitors of oxaliplatin toxic effects on the DRG. Nevertheless, in our experimental framework, we observed the significant potential of the DRG outgrowth assay as an evaluative tool for assessing further mechanistic aspects of peripheral neuropathy and treatment impacts.

### Oxaliplatin induces ROS burst that is suppressed by TXNIP inhibitor

Despite many studies conducted in cell culture to demonstrate the ROS production after treatment with oxaliplatin (18), direct evidence of this process on the extracted DRG has not been demonstrated. To explore the oxidative stress associated with oxaliplatin we used an assay where the partially outgrowth DRG (2 days) was exposed to different concentration of oxaliplatin in the presence of an ROS sensor. Measured fluorescence from the sensor within the DRG was used to judge the level of ROS. Control DRG shows relatively low level of ROS induced fluorescence within two hours of recording. Addition of oxaliplatin to this media led to the rapid increase of fluorescence within a few minutes and continued to grow for at least two hours (**Figure 3**). Predictably, the inhibition of Oct2 by cimetidine prevented oxaliplatin entry into the DRG cells and suppressed ROS production.

The cellular level of ROS and ROS -induced inflammation in many cell types is highly controlled by a thioredoxin system that removes intracellular ROS (65, 66). This thioredoxin system is composed of thioredoxin 1 (TRX1) and thioredoxin 2 (TRX2) proteins (collectively, TRX), thioredoxin interacting protein (TXNIP), thioredoxin reductase (TRXR) and other molecules. TRX1 is located in the cytosol and TRX2 is located in the mitochondria (67) -remove radicals via two closely located cysteine moieties (67). The TRX system removes radicals via two closely positioned cysteine-based thiol moieties that react with ROS molecules and form a disulfide (cystine) bond preventing ROS from oxidizing proteins, lipids, and DNA.

TXNIP is an endogenous inhibitor of thioredoxin. Under normal physiological conditions with low levels of ROS, TXNIP and TRX are bound through the formation of intermolecular disulfide bonds (68),schematically shown in **Figure 5**. However, not all TRX molecules in the cell are bound and inhibited by TXNIP. This ensures that there is always a pool of active TRX available to counteract elevation of ROS. When exposed to ROS, TXNIP detaches or displaced from TRX, allowing TRX to perform its antioxidant roles. In its active state, TRX’s two thiol groups neutralize ROS and restore proteins oxidized by ROS (69). Once TRX is oxidized by ROS or oxidized proteins, TRXR reduces TRX to its active form (70, 71).

**Figure 5.**
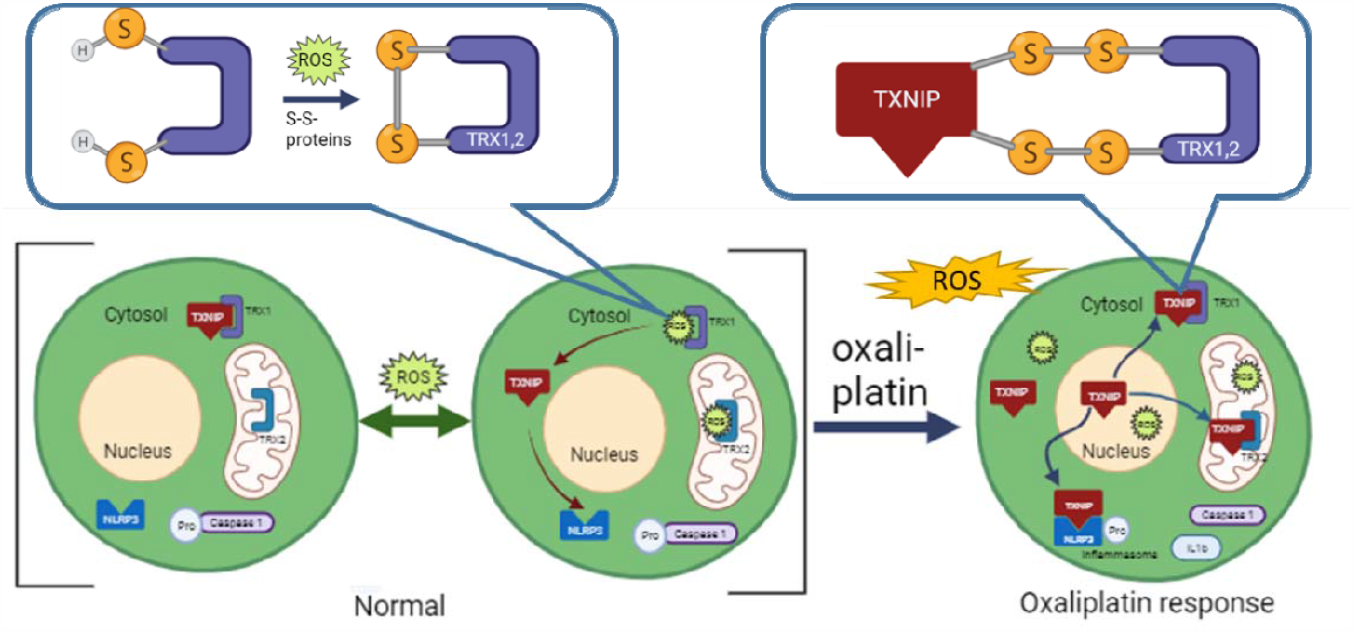
Suggested role of TXNIP in regulating ROS and inflammatory burden in DRG cells. Under normal conditions, cytosolic thioredoxin 1 (TRX1) and mitochondrial thioredoxin 2 (TRX2) suppress ROS via either direct reaction with ROS or by reducing oxidized proteins. This process is dynamically controlled by TXNIP in both cytosol and mitochondria. At higher level of stress, TXNIP is overexpressed, and TRX become less available through direct interaction between TXNIP and TRX blocking. Overproduced TXNIP shuttles to the cytosol and mitochondria where it binds TRX1 and TRX2, respectively, inhibiting the ROS elimination process. Consequently, ROS levels increase initiating mitochondrial distress. In addition, overexpressed TXNIP binds NLRP3 and triggers the formation of inflammasome NLRP3 leading to the inflammatory response and neuropathic pain.

Under a stronger level of stress, the expression of TXNIP can be upregulated leading to less active TRX activity and an increase of ROS (72). Consequently, ROS increases in the cytosol and mitochondria, initiating damage to proteins, damage to DNA, and mitochondrial distress.

In response to increased ROS levels, overexpressed TXNIP also activate the inflammatory pathway. Overexpressed binds NLRP3 and triggers the assembly of NLRP3 inflammasome complex (66) schematically shown in In response to increased ROS levels, overexpressed TXNIP can also activate the inflammatory pathway. Overexpressed TXNIP binds NLRP3 and triggers the assembly of the NLRP3 inflammasome complex (66) schematically shown in **Figure 5**, leading to the inflammation and associated cellular responses (73, 74). In neurons, the activation of the inflammasome complex generates pro-inflammatory cytokines such as IL-1β (75) which ultimately results in inflammatory neuropathic pain (76). A similar TXNIP activation mechanism has been reported in the context of diabetes induced peripheral neuropathy (77). We hypothesize that a comparable mechanism can be also responsible for the oxaliplatin-induced pain.

Given the potential connection between neuropathic pain and oxidative stress/inflammation on another, we used our DRG explant assay to investigate whether inhibiting the TXNIP’s activity with a known TXNIP inhibitor verapamil (78) suppresses the negative role of oxaliplatin on DRG regrowth and oxidative stress. Our results showed moderate but statistically significant improvement of the neurite outgrowth (**Figure 4A**) and suppression of the ROS in the DRG (**Figure 4B**) with even a small level of verapamil (ratio verapamil to oxaliplatin 1:100). Interestingly, the increase levels of verapamil never achieved the full regrowth capacity, suggesting the regrowth is partially decoupled from the TXNIP mediated ROS suppression pathway and thus more is more specific toward the oxidation stress than OCT2.

Consistent with recent literature, the observed reduction in ROS supports the link between TXNIP overexpression and the ROS-induced inflammation contributing to conditions like diabetic neuropathy (37, 79), rheumatoid arthritis (80) (73), and chronic pain from partial sciatic nerve ligation in mice (81). In humans, elevated TXNIP levels have been associated with axonal damage seen in type 2 diabetes mellitus (82) and are notably higher in the neurons of older individuals (83).

## CONCLUSIONS AND FUTURE WORK

Our work underscores the potential of the DRG explant model as an important tool in deciphering the mechanism of CIPN and testing the potential treatment. The elevation of ROS, instigated by the platinum component of oxaliplatin, not only impairs the regular functioning of antioxidant enzymes but also inflicts damage to the proteins, DNA, and mitochondria, all of which play a part in the onset and progression of CIPN. Disappointingly, most of tested antioxidants have shown only limited improvements and have not been successful in clinical trials. This amplifies the need for the identification of innovative therapeutic targets and treatments that inhibit ROS production in DRG.

In our research, we effectively showcased the DRG explant model as a robust method for examining oxidative stress mechanisms intrinsic to CIPN. Our data reveals sustained ROS elevation in DRGs exposed to oxaliplatin. Through quantitative analyses of neurite regrowth areas and oxidative stress levels, we gauged the toxic impacts of oxaliplatin and the potency of potential interventions. With its multifaceted cellular makeup and its preservation of extracellular structures, the model provides a physiologically relevant platform for screening treatments targeting both neuron and non-neuron cells in the DRG. This assay emerged as an effective model for corroborating potential CIPN targets, paving the way for future investigations toward possible therapeutic interventions.

From an experimental standpoint, the use of extracted DRG vs mice brings a superior control over the local milieu. It facilitates precise adjustments to factors like drug concentrations – a feat hard to achieve in in vivo settings. Furthermore, state-of-the-art imaging techniques can be seamlessly incorporated with the DRG. Given that a mouse harbors over 31 DRG pairs, the model is particularly suitable for a relatively (compared to live mice) high-throughput drug screenings for CIPN, expediting research while reducing the use of live animals. However, a limitation to consider is that ex vivo DRG models might overlook certain systemic elements, such as circulatory or immunological interactions, that could be substantial factors in CIPN manifestations. Despite the convenience and speed of the DRG model, ultimately we will need to proceed to animals studies prior to clinical trials. Looking forward, our focus will be to assess the potential benefits of agents like verapamil and potentially other TXNIP inhibitors in the animal models of CIPN.

## ACKNOWLEDGMENTS

M. B. acknowledges support from the NIH (R01 CA208623 and R21CA269099). Imaging results were performed in part using Washington University Center for Cellular Imaging (WUCCI) supported by Washington University School of Medicine, The Children’s Discovery Institute of Washington University and St. Louis Children’s Hospital (CDI-CORE-2015-505 and CDI-CORE-2019-813) and the Foundation for Barnes-Jewish Hospital (3770 and 4642). Imaging results were also performed in the Optical Spectroscopy Core (Washington University)

