## Supplemental File for "DRG Explant Model: Elucidating Mechanisms of Oxaliplatin-Induced Peripheral Neuropathy and Identifying Potential Therapeutic Targets"

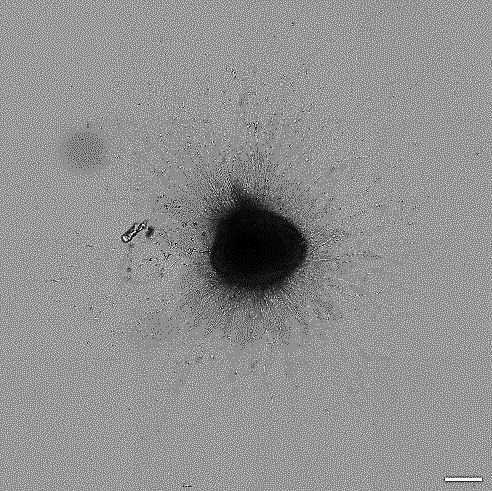

**Figure S1** Representative Bright-Field image of control DRG. Due to low contrast between background and small fibers neurites are difficult to quantify under bright field mode.

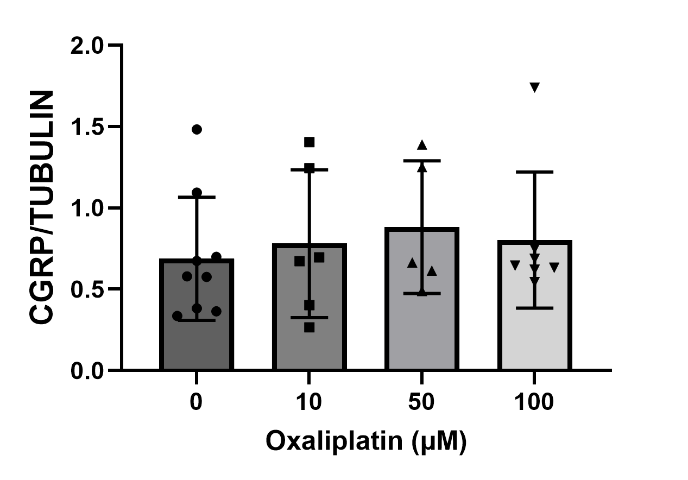

**Figure S2** Ratio of CGRP(+) neurons to total neurons stained against β-III tubulin in the intact DRG grown in the presence of different concentrations of oxaliplatin for 3 days suggest oxaliplatin is indiscriminative toward neurons in the DRG.

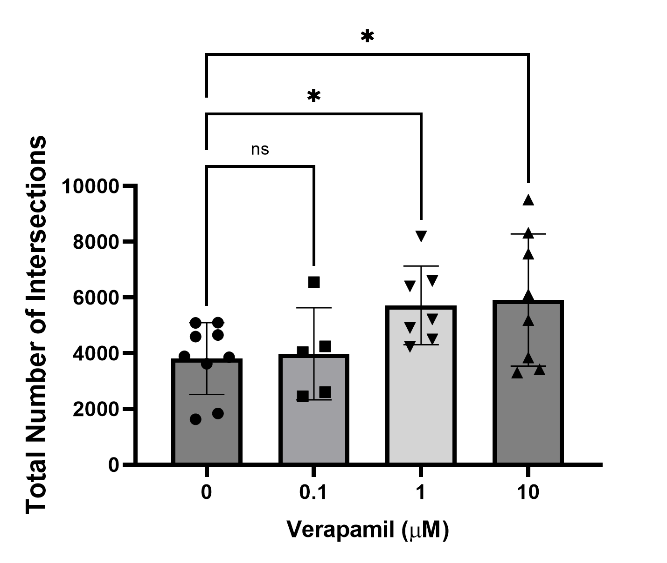

**Figure S3** **Dose-response of verapamil on the DRG regrowth.** Total number of intersections of DRG explants treated with 100 μM oxaliplatin and different dosage of verapamil. Data are expressed as the mean ± SD. *P<0.05
